# Do Dams Also Stop Frogs? Assessing Population Connectivity of Coastal Tailed Frogs (*Ascaphus truei*) in the North Cascades National Park Service Complex

**DOI:** 10.1101/062844

**Authors:** Jared A. Grummer, Adam D. Leaché

## Abstract

We investigated the effects of three hydroelectric dams and their associated lakes on the population structure and connectivity of the coastal tailed frog, Ascaphus truei, in the North Cascades National Park Service Complex. Three dams were erected on the Skagit River in northern-central Washington state between 1924 and 1953 and subsequently changed the natural shape and movement of the Skagit River and its tributaries. We collected 183 individuals from 13 tributaries and generated a dataset of >2,500 loci (unlinked SNPs) using double digestion restriction site-associated DNA sequencing (ddRADseq). An analysis of molecular variance (AMOVA) identified ~99% of the genetic variation within groups, and the remaining variation among groups separated by dams, or the Skagit River. All populations exhibited low F_*ST*_ values with a maximum of 0.03474. A ‘de novo’ discriminant analysis of principal components revealed two populations with no geographic cohesiveness. However, testing groups that were partitioned *a priori* by the dams revealed distinctiveness of populations down-river of the lowest dam. Coalescent-based analyses of recent migration suggest that up to 17.3% of each population is composed of migrants from other populations, and an estimation of effective migration rates revealed high levels of migration heterogeneity and population connectivity in this area. Our results suggest that although the populations down-river from the lowest dam are distinguishable, a high level of *A*. *truei* population connectivity exists throughout the North Cascades National Park Service Complex.

---

“I went out in my alpine yard and there it was…hundreds of miles of pure snow-covered rocks and virgin lakes and high timber…Below, instead of the world, I saw a sea of marshmallow clouds.” – Jack Kerouac on the North Cascades National Park, taken from <u>The Dharma Bums</u>

## Introduction

Identifying patterns of genetic diversity across the landscape can help mitigate negative effects brought about by human land-use and climate change on wildlife populations while also accurately guiding conservation efforts (Dudley et al. 2005; Parmesan 2006). A variety of factors can cause populations to diverge genetically, including human-mediated landscape alterations (Pess et al. 2008; Sepulveda & Lowe 2009) and natural landscape features (Spear et al. 2005). In the Pacific Northwest, river dams and heterogeneous topography have led to population differentiation in both amphibian and fish species (Funk et al. 2005; Pess et al. 2008).

Habitat fragmentation, particularly that which is largely the result of human activities, is a major cause of the worldwide loss of alpha diversity (Fahrig 2003). It can also drive population genetic processes such as increased inbreeding, loss of heterozygosity, and genetic drift, all of which can increase the probability of local extinction (Curtis & Taylor 2003). Specifically, in the Pacific Northwest, urban development and timber harvesting are two major factors that have contributed to habitat loss and fragmentation (Murphy & Hall 1981). The damming of rivers, which is both a form of urban development and habitat fragmentation, has also greatly affected organismal populations in the Pacific Northwest. The dams of the Elwha River on Washington’s Olympic Peninsula, for instance, were in place for nearly a century until their removal in 2012, and these dams have had severe impacts on the migration and habitat of fish and other aquatic species (Duda et al. 2008; Pess et al. 2008).

Washington state has nearly 1,200 dams across its 39 counties, 98 of which are hydroelectric. The rapidly growing human population is putting increases stresses on freshwater supplies across the globe. Across the U.S., more than 5 million kilometers of streams and rivers are dammed, with approximately 0.25% having any type of protection (Benke 1990; Pringle et al. 2000). Unfortunately, the damming of riverine systems often has large negative consequences for the ecosystems in those areas (Richter et al. 2003). Some ecological consequences include population reduction and extirpation of migratory fish, range fragmentation, and increases in exotic species (Pringle et al. 2000). Dams also affect terrestrial/riparian habitats and the species that occupy them by altering water levels and water availability in the areas both upstream and downstream of dams (Nilsson & Berggren 2000). Though poorly studied in this context, amphibians therefore have the potential to provide insight into both the aquatic and terrestrial effects precipitated by damming riverine systems.

The Skagit River Hydroelectric Project, which is owned by the Seattle City Light public utility company, is a hydroelectric system spanning two counties (Skagit and Whatcom) in Washington State and is composed of three dams along the Skagit River. The Skagit River courses for about 200 kilometers in the U.S., and the dams are located approximately 150 km upriver from its mouth. All three dams, and their associated lakes, are located within the North Cascades National Park (NCNP). Hydroelectric dams are a critical resource for Seattle, as they supply the city with ~88.9% of its total electricity needs (http://www.seattle.gov). Out of the total %89% of hydroelectric power that Seattle receives from all hydroelectric dams, the Skagit River dams provide ~20%. The Gorge, Diablo, and Ross dams were erected in 1924, 1930, and 1953, respectively. Through efforts to detail population genetics of the Cascades frog (*Rana cascadae*), Monsen & Blouin (2004) sampled populations from the Skagit River watershed. Their research showed that strong genetic subdivision and low migration exist at small geographic distances for this species (Monsen & Blouin 2003). The biotic effects of the Skagit River Hydroelectric Project were also assessed in lake populations of the long-toed salamander (*Ambystoma macrodactylum*). Results from this study indicated that populations west of the orographic divide (Skagit drainage) show higher levels of genetic diversity than those to the east (Stehekin drainage; Shields & Liss 2003).

Given its abundance in the NCNP and potential for long-distance dispersal, we chose to focus on the coastal tailed frog, *Ascaphus truei*, for this study. This species, along with its congener *A. montanus* (Rocky Mountain tailed frog), have a distribution limited to the mesic areas of the Pacific Northwest (loosely defined as the area from northern California to British Columbia, and eastward into western Idaho). *Ascaphus truei* individuals can be locally abundant in first-and second-order streams, however, their restrictive physiology requirements for both lower temperatures and high moisture levels have been thought to lead to limited dispersal (Claussen 1973; Brown 1975). In a mark-recapture study, Wahbe et al. (2004) captured a majority of *A. truei* individuals within 50m of streams, with a small number of individuals discovered between 50-100m from the stream. Notwithstanding, Corn & Bury (1989) documented *A. truei* individuals as far xsas 1km from the closest stream. The results of the Corn & Bury (1989) study are in-line with more current research, which has uncovered high levels of gene flow at relatively large distances and population connectivity of *A. truei* populations at a scale of 25-30km on Washington’s Olympic Peninsula (Spear & Storfer 2008).

We assessed genetic connectivity and structure using single nucleotide polymorphisms (SNPs). A variety of methods have been developed in the past few years to acquire SNPs from across the genome of non-model organisms (e.g. Elshire et al. 2011; Etter et al. 2011; Peterson et al. 2012). These methods are capable of producing datasets consisting of thousands of unlinked loci from across the genome. Furthermore, these methods are appealing because they require little knowledge of the genome, which eliminates the need to invest in genomic resource development. The large numbers of loci that these methods produce have the ability to provide accurate estimates of important population genetic parameters and increased statistical power for estimating population differentiation (Felsenstein 2006; Bj00F6;rklund & Bergek 2009). These datasets hold particular promise for identifying population structure caused by processes that have occurred in recent timescales.

In this study, we aimed to determine whether or not anthropogenic alterations to the landscape, hydroelectric dams and concomitant lakes in this case, affect population connectivity of *A. truei* in the NCNP. Because the Skagit River courses through a narrow canyon in our study site, dams have raised the water level some 400’ above the “natural” elevation of the river, which not only separates streams from opposite sides of the river that used to be much closer, but also places stagnant water in between suitable *A. truei* habitat. This is one of the reasons why we expect to see population structure in this species caused by these dams. We collected nearly 200 individuals over two years from thirteen named streams and their associated tributaries in areas around these dams and their associated lakes that allowed us to test three explicit hypotheses: populations of *A. truei* are structured by a) the hydroelectric dams, b) the Skagit River and the lakes created by the dams, or c) a combination of the dams and the Skagit River. Our null hypothesis is that no population structure exists in this area. To address these hypotheses, we constructed a dataset consisting of ~2,500 SNPs to determine both population structure and genetic connectivity amongst sampled populations.

## Materials and Methods

### Sample Collection

We collected 196 *A. truei* individuals between 2012-’13 from 13 streams and their tributaries in the NCNP, comprising 25 unique localities (Fig. 1; under U.S. Department of the Interior and Washington Department of Fish and Wildlife permit nos. N0CA-2012-SCI-0044, N0CA-2013-SCI-0013, and RCW 77-32-240, WAC 220-20-045). Seven individuals were adults, 189 were larvae, and all that were preserved as vouchers were deposited into the University of Washington’s Burke Museum Herpetology and Genetic Resources Collections (see Supplemental Table 1 for UWBM accession numbers and locality information).

**Figure 1:**
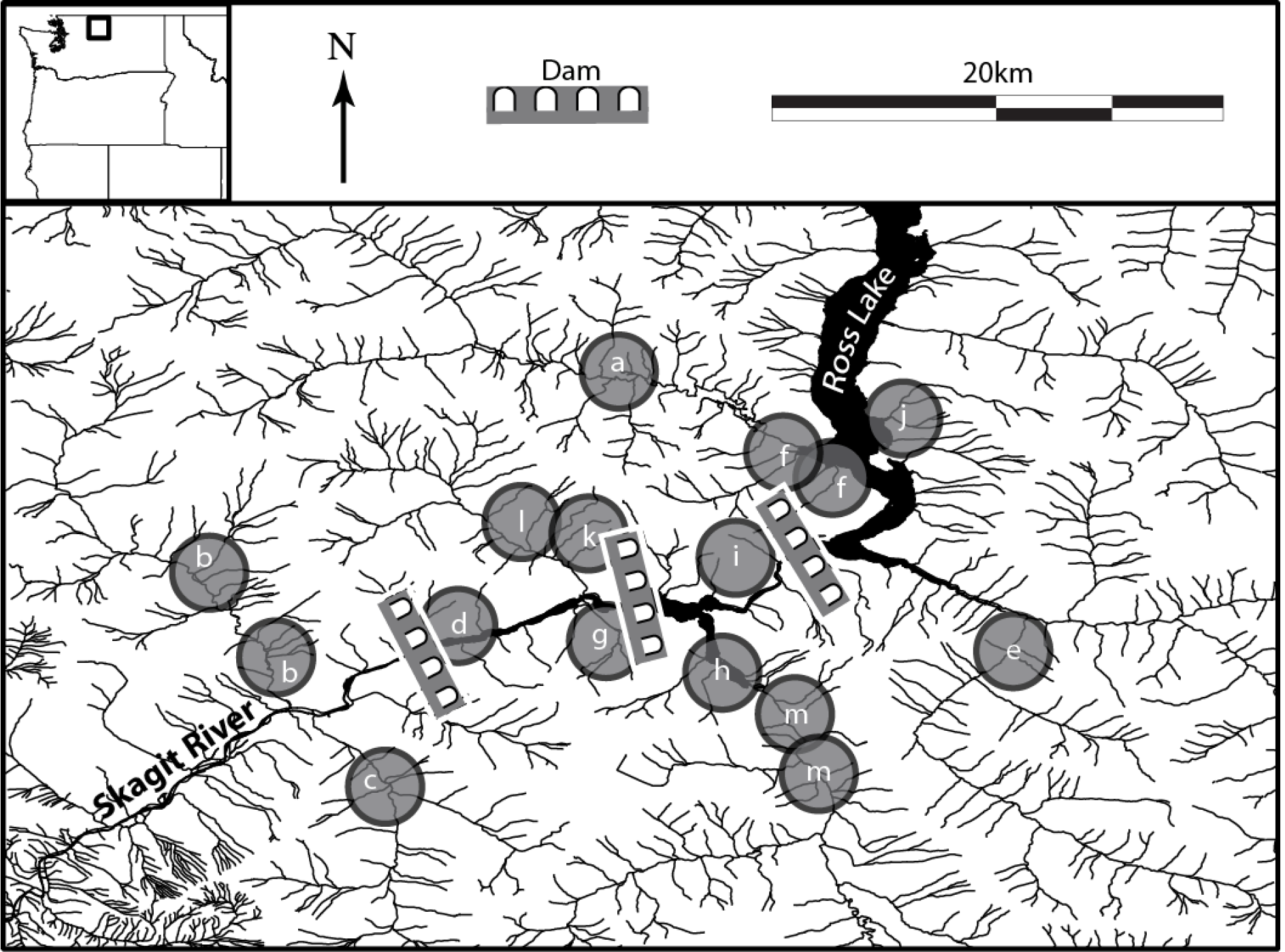
Study area showing sampling sites and waterways in the North Cascades National Park Service Complex. Letters correspond to the following streams, with sample sizes following stream name: a – Big Beaver (16), b – Goodell (15), c – Newhalem (15), d – North Gorge (15), e – Panther (13), f – Pierce (14), g – Pyramid (20), h – Rhodes (15), i – Riprap(13), j – Roland (13), k – Sourdough (7), l – Stetattle (14), m – Thunder (13). Refer to supplementary table S1 for further sampling information

**Table 1:** Population structure hypotheses tested in our analyses. Total number of groups tested under each hypothesis is shown, where numbers indicate group assignment that correspond to Fig. 2.

| Hypothesis | # Groups | Big<br>Beaver | Panther | Pierce | Roland | Rhodes | Riprap | Thunder | North<br>Gorge | Pyramid | Sourdough | Stetattle | Goodell | Newhalem |
| --- | --- | --- | --- | --- | --- | --- | --- | --- | --- | --- | --- | --- | --- | --- |
| By Dam | 4 | 1 | 1 | 1 | 1 | 2 | 2 | 2 | 3 | 3 | 3 | 3 | 4 | 4 |
| By River | 2 | 1 | 2 | 1 | 2 | 2 | 1 | 2 | 1 | 2 | 1 | 1 | 1 | 2 |
| By Dams<br>and River | 8 | 7 | 8 | 7 | 8 | 6 | 5 | 6 | 3 | 4 | 3 | 3 | 1 | 2 |

### DNA Data Collection

Genomic DNA (gDNA) was extracted from either liver, toe clip, or tail clip using the Qiagen DNeasy extraction kit (Qiagen, Valencia, CA). Total gDNA quality was assessed qualitatively through visualization on a 1% agarose gel and quantitatively with a Qubit fluorometer (Life Technologies, Carlsbad, CA). Thirteen samples were discarded from analyses due to poor gDNA and/or data quality. We generated sequence data using the double digestion restriction site-associated DNA sequencing (ddRADseq) technique developed by Peterson et al. (2012). Samples were first digested for eight hours at 37°C with the restriction enzymes SbfI(“rare” 8bp restriction site sequence [5’ CCTGCAGG 3’]; New England Biolabs, Ipswich, MA) and MspI (“common” 4bp restriction site sequence [5’ CCGG 3’]; New England Biolabs). The enzyme T4 DNA ligase (New England Biolabs) was then used to ligate barcoded oligonucleotides to each genomic DNA fragment (each barcode 5bp in length) that were unique to each row of individuals on the sequencing plate. Individuals were then pooled, followed by a size selection step with the Blue Pippin (Sage Science, Beverly, MA) where all loci between 415-515bp were retained. A final PCR step using Phusion Taq polymerase (New England Biolabs) with the following thermocycler conditions was conducted to amplify all loci and attach a 6bp index unique to each pool for sequence de-multiplexing: 98°for 0:30, (98°for 0:10, 58°for 0:30, 72°for 0:30) × 12 cycles, and a final 10:00 extension at 72°C. Ninety-six or 144 individuals were multiplexed across two separate sequencing runs at the University of California Berkeley QB3 Vincent J. Coates Genomics Sequencing Laboratory on an Illumina HiSeq 2500 with 50bp single-end sequencing.

### DNA Data Assembly

Raw Illumina reads were processed with the program pyRAD v3.0.5 (Eaton 2014) to generate alignments of phased SNPs (single nucleotide polymorphisms). Reads were discarded if they had ≥4bp with a Phred quality score <20. Samples were first de-multiplexed based on unique barcode-index combinations, then sequence “clusters” were generated by pyRAD using the programs VSEARCH (https://github.com/torognes/vsearch) and MUSCLE (Edgar 2004). Reads were first clustered within individuals into loci that were ≥90% similar, and then across individuals with the same threshold. Loci were retained if they had a minimum sequencing depth of 10x. PyRAD also applies a paralog filter in which the user specifies the threshold value, which represents the maximum percentage of individuals allowed to have a heterozygous base (IUPAC “ambiguities”) at a given site. A higher value for the paralog filter results in more heterozygotes at any given position because of (a) fixed allelic differences, or (b) sequence polymorphism, both of which can appear the same due to sequence reads containing both alleles. For our final datasets, we set this value fairly high at 90%, meaning ≤90% of the individuals at a given locus (unlinked SNP) could share a sequence heterozygosity, because we expect that heterozygosity can occur at a high frequency at this limited spatial scale. Finally, we compiled two final datasets that differed with respect to amount of missing allowed: one consisted of a missing data level of 50%, meaning that ≤50% of the individuals can have missing data (“?”) at a given SNP, whereas the second dataset contained 0% missing data, meaning every individual had data for every locus.

### Identifying Genetic Subdivision

After data quality control and assembly, our final datasets consisted of 183 individuals and 2,537 and 211 unlinked SNPs (loci) for the 50% and 100% complete datasets, respectively. We aimed to test three *a priori* hypotheses (along with the null hypothesis of no genetic structure) to determine which geographical feature (including dams), if any, is responsible for causing genetic subdivision between *A. truei* populations: genetic subdivision caused by a) three hydroelectric dams, b) the Skagit River, or c) a combination of the dams and the Skagit River (Table 1).

We first tested for a correlation between genetic and geographic distances (isolation by distance) with a Mantel test in the program Adegenet (Jombart 2008; Jombart et al. 2010; Jombart & Ahmed 2011). The significance of isolation by distance was tested by creating a null distribution (an absence of spatial structure; 1000 replicates) and comparing the empirical value to this distribution. We next assessed genetic variation by these pre-defined groups using an analysis of molecular variance (AMOVA) in Arlequin (v3.5; Excoffier & Lischer 2010), where a locus-by-locus AMOVA was performed on the 100% complete dataset (results from all loci were combined for the final result). We also used Arlequin to calculate population-pairwise F_*ST*_ values, which were done with 1,000 permutations to test for statistical significance of population differentiation.

We identified the number of populations (*k*) using a discriminant analysis of principal components (DAPC) on the 100% complete dataset in the R package Adegenet (Jombart 2008; Jombart et al. 2010; Jombart & Ahmed 2011). The program first transforms the SNP dataset through a principal component analysis (PCA), then a discriminant analysis (DA) is performed on the output of the PCA analysis. Generally, the DAPC method seeks to maximize between-group genetic variation while minimizing within-group variation, and has the benefit of DAPC over other population clustering methods in that it makes no assumptions of the underlying population genetic model. We first performed a “*de novo*” analysis without individuals assigned to populations, and we chose the optimal *k* value based on the Bayesian Information Criterion (BIC) of the likelihood score associated with each *k* iteration (“find.clusters” command). We then assigned individuals to populations based on the groups defined in Table 1 and explored population structuring under these three hypotheses. For all analyses, we used 60 principal components, which is approximately one-third the number of individuals and the recommended amount by the program authors.

We also used the program Admixture (Alexander et al. 2009) to determine *k* This program is similar to the more popular program Structure (Pritchard et al. 2000) in that both approaches model the probability of the observed genotypes using ancestry proportions and population allele frequencies. However, one difference is that unlike Structure, which utilizes a Bayesian algorithm, Admixture uses a maximum likelihood approach that differs in how the optimal *k* value is selected. Specifically, the Evanno et al. (2005) method widely used in Structure analyses cannot evaluate *k*=1. In contrast, the cross-validation method employed in Admixture can evaluate *k*=1. We ran Admixture (Alexander et al. 2009) on our 50% complete dataset, running 10 replicate analyses with unique starting seeds to ensure consistency of results.

### Estimating Migration Rates

The human-mediated habitat change that we assessed in this study was very recent, within the past ~70 years. This amount of time equates to approximately 8-40 generations, given an estimated generation time of 2-8 years for *A. truei* (Bury & Adams 1999; Nielson et al. 2001). To estimate recent migration rates over the last several generations, we used the Bayesian program BayesAss v3.0.4 (Wilson & Rannala 2003). This program estimates the proportion of each population that are immigrants from each of the other populations (e.g., asymmetric migration rates), in addition to estimating the total number of nonimmigrants, and first-and second-generation migrants. A benefit of this program over others that estimate migration is that it relaxes many population genetic assumptions such that the populations do not have to be at equilibrium. Importantly, genotype frequencies can deviate from Hardy-Weinberg equilibrium within populations. We ran four replicate analyses on our 50% complete dataset, which was partitioned in four different ways: three analyses where individuals were assigned to groups based on Table 1, and the fourth where individuals were assigned to the stream in which they were sampled. Each was run for 10^8^ generations, and the first 2×10^7^ generations were discarded as burnin (with “mixing” parameters-a 0.4-f 0.1-m 0.2). Convergence was visually assessed in Tracer v1.5 (Rambaut & Drummond 2007).

Secondly, we estimated migration rates using the “estimating effective migration surfaces” (EEMS) method (Petkova et al. 2015). This method is based upon a stepping stone model in which migration is allowed between neighboring demes in a grid, the density of which the user specifies. This approach assesses genetic connectivity across the landscape in a way that makes it conceptually related to methods that utilize circuit theory (Hanks & Hooten 2013). Migration rates are adjusted such that the genetic differences expected under the model are close to the genetic differences observed in the data. These estimates are then interpolated across the landscape to produce the estimated effected migration surface. This method requires *n*sites *≫n* individuals, so we used the larger (50% complete) data matrix that was 2,341 loci after removing non-biallelic sites. We experimented with the number of demes (grid density) and ultimately used 100 demes, and ran the analysis for 5 × 10^7^ generations with 10^7^ generations as burnin and saving the chain state every 50,000 generations. Convergence was assessed by concordance across replicate runs with different starting seeds, in addition to examining the trace plot of the MCMC posterior.

## Results

### Population and Genetic Structure

Our final datasets consisted of 183 individuals and 2,537 or 211 loci (SNPs) for the 50% and 100% complete matrices, respectively; the number of individuals per stream ranged from 7-20 (Fig. 1; Table S1). We did not detect any significant signal of isolation by distance in our dataset (*p* = 0.10; Supplemental Fig. 1). Our AMOVA results revealed that the vast majority of genetic variation is found within groups (>99%), e.g., not between pre-defined groups (Table 2). Whether the data were divided into groups separated by the dams, the Skagit River, or the dams and Skagit River made no significant difference in among-group genetic variation, which was quite low in all cases at <1% (Table 2).

**Table 2:** Results from AMOVA analyses on the 100% complete dataset. Refer to Table 1 for assignments of streams to groups.

| Source of Variation | Percentage of Variation |  |  |
| --- | --- | --- | --- |
|  | By Dams | By River | By Dams and River |
| Among Groups | 0.82 | 0.16 | 1.00 |
| Within Groups | 99.18 | 99.84 | 99.00 |

Estimates of F_*ST*_ between groups ranged from 0.00 to 0.03474 (Tables 3–5), indicating little differentiation between *a priori* defined groups. However, in spite of these low values, the majority of pairwise population differentiation tests were significant at p <0.05 (Tables 3–5).

**Table 3:** Pairwise F_*ST*_ results when individuals were partitioned by dams and the Skagit River. Bold values indicate significance at p <0.05; all other values are insignificant.

|  | Dam Section |  |  |  |
| --- | --- | --- | --- | --- |
|  | North of Ross | Ross to Diablo | Diablo to Gorge | South of Gorge |
| North of Ross | 0 |  |  |  |
| Ross to Diablo | **0.00914** | 0 |  |  |
| Diablo to Gorge | **0.00768** | 0.00217 | 0 |  |
| South of Gorge | **0.01882** | **0.00893** | **0.01207** | 0 |

**Table 4:** Pairwise F_*ST*_ results when individuals were partitioned by the Skagit River. The single bold value indicates significance at p <0.05.

|  | Side of River |  |
| --- | --- | --- |
|  | North/West of Skagit | South/East of Skagit |
| North/West of Skagit | 0 |  |
| South/East of Skagit | **0.00192** | 0 |

**Table 5:** Pairwise F_*ST*_ results when individuals were partitioned by dams and the Skagit River. Bold values indicate significance at p <0.05; all other values are insignificant.

|  | Dam and River Partition |  |  |  |  |  |  |  |
| --- | --- | --- | --- | --- | --- | --- | --- | --- |
|  | North of Ross<br>West of Skagit | North of Ross<br>East of Skagit | Ross-Diablo<br>West Skagit | Ross-Diablo<br>East of Skagit | Diablo-Gorge<br>North of Skagit | Diablo-Gorge<br>South of Skagit | West of Gorge<br>North of Skagit | West of Gorge<br>South of Skagit |
| N Ross, W Skagit | 0 | 0 |  |  |  |  |  |  |
| N Ross, E Skagit | **0.00857** | 0 |  |  |  |  |  |  |
| Ross-Diablo, W Skagit | **0.01904** | 0.00 | 0 |  |  |  |  |  |
| Ross-Diablo, E Skagit | **0.01488** | **0.00755** | 0.00603 | 0 |  |  |  |  |
| Diablo-Gorge, N Skagit | **0.01877** | **0.00806** | 0.00261 | **0.00674** | 0 |  |  |  |
| Diablo-Gorge, S Skagit | **0.00757** | 0.00529 | 0.00195 | 0.00 | 0.00036 | 0 |  |  |
| W Gorge, N Skagit | **0.02633** | **0.03474** | **0.03057** | **0.02326** | **0.02048** | **0.01255** | 0 |  |
| W Gorge, S Skagit | **0.02203** | **0.01906** | 0.01247 | 0.00878 | **0.01334** | 0.00061 | **0.0159** | 0 |

The results from our DAPC analyses are shown in Figure 2. Without individuals assigned to populations (“de *novo*”), *k*=2 was selected with the BIC (Supplemental Fig. 2). It is interesting to note that when using the 50% complete dataset (2,537 loci), *k*=4 was the best grouping based on BIC score (though only slightly better than *k*=3; Supplemental Fig. 3). There appears to be no geographic structure with the two populations identified in the *de novo* analysis (Fig. 2a), nor with the three or four populations identified with the 50% complete dataset (Supplemental Fig. 4).

**Figure 2:**
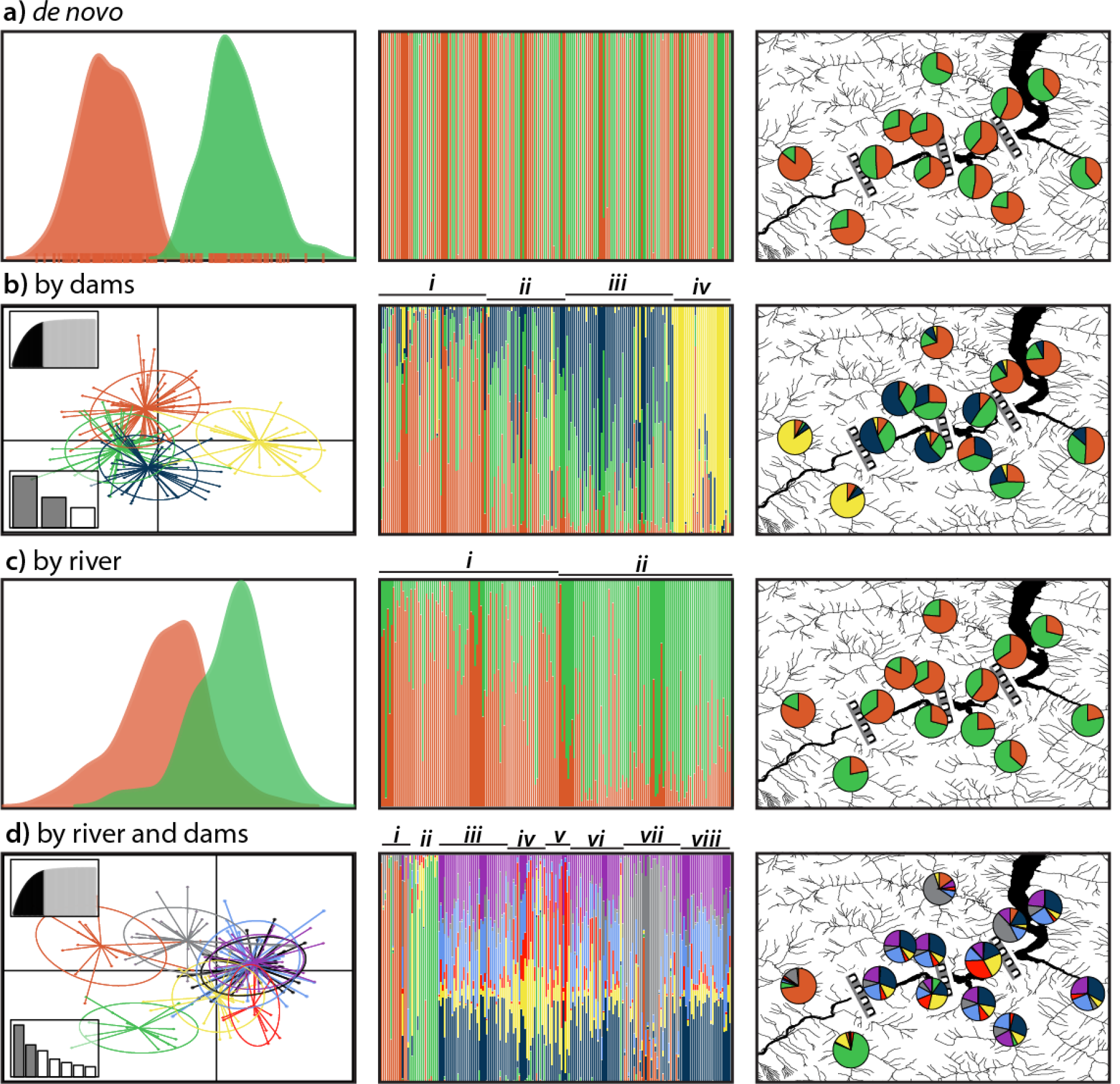
Discriminant analysis of principal components (DAPC) results from Adegenet under three different hypotheses of genetic subdivision along with *de novo* clustering. The left-hand column shows the DAPC plots, middle column shows the population assignments of individuals to each respective cluster, and the right column shows the percentage of individuals from each stream assigned to each cluster. Insets in the left column of rows (b) and (d) indicate number axes retained in principal component (top-left) and discriminant analyses (bottom-left). Roman numerals above figures in the middle column correspond with population assignments in Table 1

**Figure 3:**
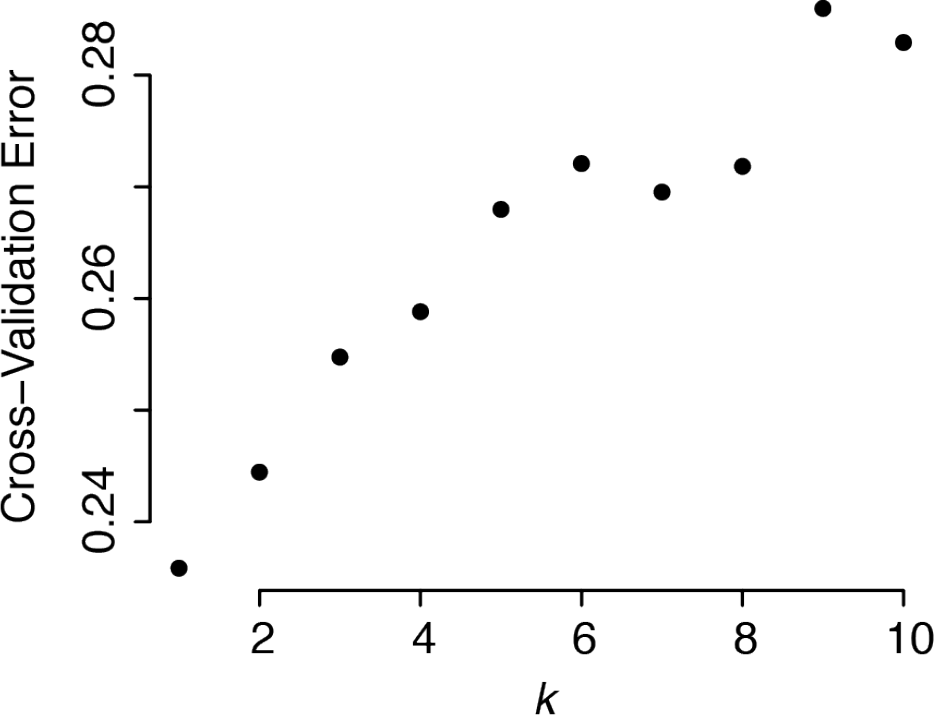
Results from the Admixture cross-validation test. The *k* value with the lowest cross-validation error is the most likely number of populations

When partitioned by dam (Fig. 2b), populations below Gorge Dam appear distinct. The populations above Ross Dam are also distinct, though with a fair amount of genetic similarities to central populations (Fig. 2b). Central populations (Ross to Diablo and Diablo to Gorge Dam stretches) are more admixed than northeastern and southwestern populations above Ross Dam and below Gorge Dam, respectively (Fig. 2b). Partitioning individuals by the Skagit River provided moderate genetic differentiation between these two clusters (Fig. 2c). And lastly, partitioning individuals by regions isolated by both dams and the Skagit River resulted in clear differentiation of the populations west of the Gorge Dam and both north and south of the Skagit River (Fig. 2d). Individuals from the Big Beaver drainage north of Ross Dam and West of the Skagit River also showed some distinctiveness from the other groups.

We also evaluated the ability of the data in its effectiveness to “correctly” assign individuals to their *a priori* defined groups. Though a characteristic of the data and its informativeness, this is largely a function of the congruence of the actual population structuring (as seen in the genetic data) with our pre-defined groups. We considered individuals to be “correctly” assigned if their assignment probabilities were >0.5 to their predefined group. The population partition that had the highest number of correctly assignments was the population composed of individuals from Goodell and Newhalem Creeks, below Gorge Dam; 90% of the individuals were correctly assigned to this group (Table 6). These two creeks also had high levels of correct assignments when partitioned by both dam and river (75% and 87%, respectively). However, when examining population structure putatively structured by both dam and river, three populations had 0 individuals correctly assigned. When considering each population structuring hypothesis, populations structured by the Skagit River had the highest percentages of correctly assigned individuals at 85% for the population north of the river and 81% for the population south of the river.

**Table 6:** Percent of individuals correctly assigned to their *a priori* defined groups during DAPC analysis for each hypothesis of population structuring. An individual is “correctly” assigned to a group when ~50% of its inferred assignment probability is to the group it was assigned to before analysis. Group numbers in the left column refer to group assignments in Table 1.

|  | Percent Correctly Assigned |  |  |
| --- | --- | --- | --- |
|  | By Dams | By River | By Dams and River |
| Group 1 | 62 | 85 | 75 |
| Group 2 | 40 | 81 | 87 |
| Group 3 | 63 | — | 0 |
| Group 4 | 90 | — | 5 |
| Group 5 | — | — | 23 |
| Group 6 | — | — | 0 |
| Group 7 | — | — | 60 |
| Group 8 | — | — | 0 |

Our Admixture (Alexander et al. 2009) analysis identified the most likely number of populations as *k*=1. These results were stable across all 10 replicate runs.

### Migration Rates

For our BayesAss results, variation in the posterior mean migration estimates across four replicates of each geographic partitioning scheme (i.e., hypotheses in Table 1) was very low (often <0.000*x*), in spite of low effective sample size (ESS) values for the overall log-probability of each analysis (results not shown). Standard deviation of the posterior mean migration estimates were also low (<0.06), so the migration estimates we present here for each hypothesis are from a single analysis. Migration rates across all geographic partitioning schemes were relatively high (>0.0075, Supplemental Tables 2-5; note that values are expressed as proportion of the population, not *N_e_m*).

The highest migration rate we observed was from the population north and west of the Skagit River into the population on the opposite side of the river (%17%; Supplemental Table 4), whereas overall the lowest migration rates were those between populations partitioned by both dam and river (lowest value of 0.76%; Supplemental Table 5).

The results of our effective migration rates analysis indicated a high level of migration heterogeneity in this area. The lower Skagit River and Gorge Dam appear to be strong barriers between populations sampled in that area (Fig. 4). In contrast, a high level of migration is inferred between populations immediately above and below Ross Dam (Pierce and Riprap, respectively; Fig. 4). Similarly, there appears to be population connectivity across both the Skagit River and Diablo Dam between the Pyramid, Rhodes, and Sourdough populations in the central portion of the study area.

**Figure 4:**
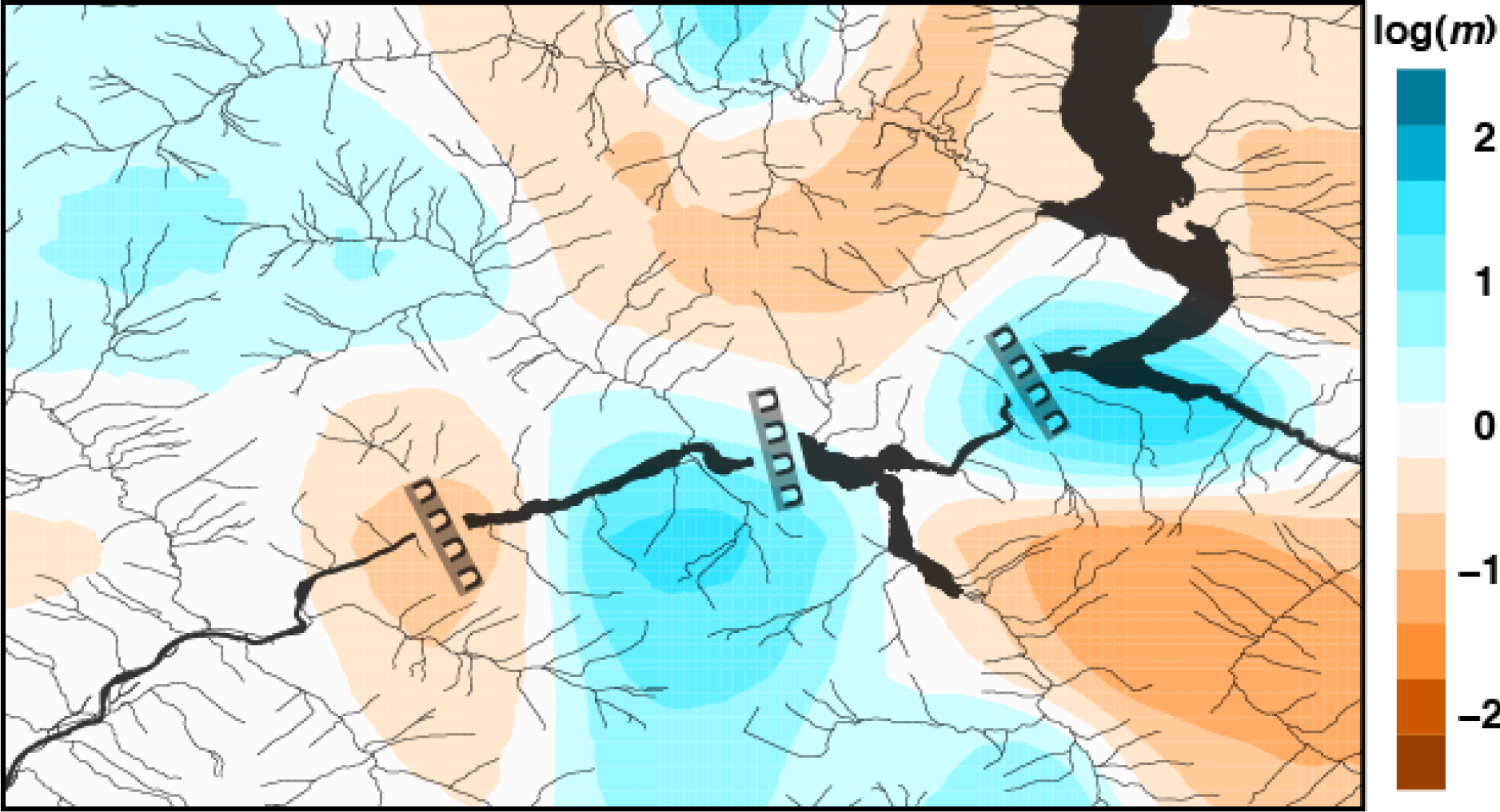
EEMS results showing estimated effective migration rates in the North Cascades National Park Service Complex. Darker blues indicate high estimated migration rates, whereas dark orange indicates low migration rates, relative to the overall migration rate across the entire area. Thus, in the log scale on the right, a value of 2 corresponds to effective migration rates that are 100× higher than the average rate

## Discussion

Even though the Skagit River Hydroelectric Project was completed between the years of 1921 and 1953, the North Cascades National Park was not established until 1968. And in spite of the fact that *A. truei* is listed as a “species of concern” by the Washington Department of Natural Resources (due to logging and other forms of habitat destruction; http://www.dnr.wa.gov), it is listed as a species of “least concern” by the IUCN (http://www.iucnredlist.org/). We found evidence for population structuring in the North Cascades National Park Service Complex associated with Gorge Dam (the furthest down-river and oldest dam) and the Skagit River/lakes (Figs. 2 and 4). This result indicates that geographic structure should be considered in future tailed frog conservation/management decisions in this area, because these populations could become more structured and/or isolated in the future. There is relatively little to no support for genetic structuring among the remaining *A. truei* populations sampled in our study.

### Ability to Detect Recent Cessation in Gene Flow

In 1921, construction of the first dam of the Skagit River Hydroelectric Project was initiated (Gorge dam); this dam is also the furthest downriver of the three. The next dam upriver, Diablo, was completed in 1930. And finally, Ross Dam, which is the furthest upriver and largest of the three (540’ tall), was completed in three stages between 1940 and 1953. Given that these dams (and therefore their associated lakes) were created between approximately 60 and 80 years ago, and a large generation time for *A. truei* of ~2-8 years (Bury & Adams 1999; Nielson et al. 2001) (which has been correlated with slow evolutionary and metabolic rates, at least for mitochondrial DNA; Martin & Palumbi 1993), ~8-40 *Ascaphus* generations have elapsed during this time; this has left little time for genome-wide mutations to accumulate due to genetic drift. The SNP dataset presented here was able to detect population structure associated with the oldest dam (Gorge Dam), but unable to detect similar structure across the more recent dams.

Our population assignment (DAPC) and effective migration rate (EEMS) analyses both indicated some level of population structuring and areas of reduced gene flow in this system. The populations that are the most differentiated from the others were sampled in Goodell and Newhalem Creeks that lie to the south and west of Gorge Dam (Figs. 1 and 2). This result could be due to the age of Gorge Dam, which is the oldest of the three (construction completed approximately 5 and 27 years before Diablo and Ross Dams, respectively). The earlier establishment time of Gorge Dam means that its presence and Gorge Lake have been putatively affecting organismal populations for more time than the other dams, albeit for a short duration before Diablo Dam and Diablo Lake. Hypothesizing that dam age is correlated with genetic differentiation of populations separated by the dams can be tested by examining F_*ST*_ (Table 3) and “correct” population assignments (Table 6), where we expect both of these values to be positively correlated with dam age. However, we do not see this pattern in our results, since the most recent dams do not fit this expectation.

Another reason for the differentiation of Goodell and Newhalem Creeks from all others is that these two populations are lower in elevation than most of the other sites we sampled (~380m above sea level), with the highest (Panther Creek) at %830m. Elevation has been shown to structure amphibian populations in other systems in northwestern North America (Giordano et al. 2007). However, high elevation sites in that study exceeded 1200m elevation, therefore the difference in elevation across our study site is likely not enough to drive the genetic differentiation we have documented by itself. And broadly, the greater structuring we see from our DAPC results may simply be due to the way the data are analyzed in this approach. This discriminant analysis seeks to maximize the (genetic) separation between groups while minimizing the variation within groups (Jombart et al. 2010). Although this method is not going to detect structure when it is not present, it will likely be more sensitive to detecting subtle differences between populations.

Though strongly favored, the *k*=2 DAPC results (and *k*=3 results for the 50% complete dataset, not shown) are difficult to interpret in a geographic context. This pattern of non-geographical population structuring could be due to two reasons. Firstly, the DAPC method lacks population-genetic assumptions that the other methods make (all of which are designed for analyzing data at the intraspecific level), perhaps causing discrepancy between this analysis and the rest. Secondly, this structure could be the result of the dams and their lakes, but viewed in the early stages of population separation and the sorting of ancestral polymorphisms through genetic drift.

### Missing Data Levels and Dataset Size

Missing data had an effect on the outcome of some of our analyses. In pyRAD (Eaton 2014), the user can modulate dataset size for a given number of individuals (in part) by changing the missing data threshold. For instance, allowing a locus (SNP) to be retained in our dataset if it has a minimum of 9/183 individuals present (%5% complete, 95% incomplete) results in a dataset size of 9,767 loci, whereas a dataset composed of only 100% complete loci (183/183 individuals at each locus) results in a dataset size of 211 loci (Supplemental Fig. 5). Thus, having more SNPs/loci comes at the cost of increasing the level of missing data. Altering the amount of missing data at a locus is not expected to change results of AMOVA or F_*ST*_ analyses, but it will increase the variance about these estimates (J. Felsenstein, pers. comm.).

We were able to see the effects of changing dataset size (and “missingness”) in our analyses. Using the 50% complete dataset in AMOVA analyses produced negative variance components (results not shown), which is difficult to interpret analytically; AMOVA analyses on the 100% complete dataset did not suffer from this problem. With DAPC analyses, *k*=3 or 4 (BIC difference of 0.15 points; Supplemental Fig. 3) is selected with the 50% missing dataset, in contrast with *k*=2 for the 100% complete dataset. The level of missing data might play a role with these differences, however, the difference is likely due to the increase in information content/loci with the higher level of missing data. We are not aware of any studies examining the effect of missing data levels on population clustering methods, but this area certainly needs to be explored.

### Conservation Implications

Natural populations are facing a variety of threats at multiple scales, from climate change (global) to habitat destruction (local). Amphibians in particular show higher modern extinction rates than those of birds or mammals, which is largely being driven by habitat loss and overutilization (Stuart et al. 2004; Hof et al. 2011). In this light, amphibians have often been portrayed as the “canary in the coal mine” to indicate the overall health and functionality of an ecosystem (Roy 2002), particularly in the Pacific Northwest (Welsh Jr & Ollivier 1998; Welsh Jr & Hodgson 2008). However, research by Kerby et al. (2010) showed that compared to a variety of taxa (fish and arthropods), amphibians are mediocre indicators of environmental health (sensitivity to contaminants in this case). The fact that coastal tailed frogs are intimately tied to riparian systems and have two distinct life stages in water and on land made them an ideal candidate to address our hypotheses in this study.

Although we did see some signs of population structuring due to the Gorge Dam and Skagit River/lakes created by the dams, there are relatively high levels of gene flow and migration across the landscape in our study site. Because of the high mortality before metamorphosis in this species, the vast majority of individuals that we sampled in this study were larvae (“tadpoles”; 178/183); three out of the five adults were non-lethally sampled and released (toeclip). Daugherty & Sheldon (1982) documented high vagility amongst younger age classes of the congener *A. montanus*, with strong philopatry amongst adults. Given the strongly larval-biased composition of our dataset, results might have revealed stronger genetic structuring by geography if we would have sampled more adults. *Ascaphus truei* populations in this area appear to be healthy and show little effect due to the presence of the Skagit River Hydroelectric Project at this time.

## Conclusions

In this study, we examined the genetic structure and population connectivity of *Ascaphus truei* in the North Cascades National Park Service Complex using a large molecular dataset composed of hundreds and thousands of loci sampled from across the genome. We specifically tested hypotheses that aimed to determine if man-made structures, in this case hydroelectric dams and the lakes that they have created, or natural landscape features (the Skagit River) were responsible for causing population structure in this species. Using genome-wide SNP data, we are able to detect population structure at a fine geographic scale that coincides with the oldest Dam (Gorge Dam), suggesting that SNP data are able to detect recent population structure that may have elapsed over as few as 12 generations from when Gorge Dam was erected. In addition to this new evidence for population structure below Gorge Dam, we found evidence for high levels of population connectivity throughout the North Cascades National Park System Complex.

## Acknowledgments

We first and foremost graciously thank the Wildlife Research Program from the Seattle City Light public utility company for providing a generous grant that enabled this research. Ron Tressler with the Seattle City Light and Becky Johnson in the University of Washington Department of Biology were also extremely helpful in coordinating logistics and finances of this project. Special thanks to M. Miller, I. Caviedes-Solis, A. Cortes, A. Gottscho, N. Herschberger, B. Peecook, H. Rockney, P. Tosello, D. Vilhena, and J. Wilkey for assistance with fieldwork. We wish to thank National Park Service personnel A. Rawhouser and B. Kuntz for aid in study design, R. Rochefort with acquisition of collecting permits, and North Cascades National Park superintendent P. Jenkins for supporting this research. Lastly, we thank B. Rannala for his assistance with BayesAss analyses. This work was facilitated though the use of advanced computational, storage, and networking infrastructure provided by the Hyak supercomputer system at the University of Washington. The completion of this work was aided by a pre-doctoral fellowship from the National Institutes of Health given to JAG.

## Data Accessibility

– DNA Sequences: Genbank accession nos. xxxx-xxxx
– All collecting locality information is available in the Supplementary Materials section.

## Author Contributions

JAG and ADL conceived and designed the project. JAG completed the fieldwork, lab work, analyses, and wrote the manuscript. ADL provided lab resources, helped with analyses, and edited the manuscript.

